# Unsupervised classification of graded animal vocalisations using fuzzy clustering

**DOI:** 10.1101/2024.09.13.612808

**Authors:** Benjamin Benti, Patrick JO Miller, Heike Vester, Florencia Noriega, Charlotte Curé

**Affiliations:** Cognitive and Social Ethology Team, University Clermont-Auvergne, Clermont-Ferrand, France; Sea Mammal Research Unit, University of Saint Andrews, Saint Andrews, United Kingdom; UMRAE Cerema-Université Gustave Eiffel, Laboratory of Strasbourg, France; Ocean Sounds e.V., Marine Mammal Research and Conservation charity, Berglen, Germany; CODE University of Applied Science, Berlin, Germany

**Keywords:** fuzzy clustering, gradation, long-finned pilot whales, MFCC, unsupervised classification

## Abstract

We present here an unsupervised procedure for the classification of graded animal vocalisations based on Mel frequency cepstral coefficients and fuzzy clustering. Cepstral coefficients compress information about the distribution of energy across the frequency spectrum into a reduced number of variables and are well-defined for signals of various acoustic characteristics (tonal, pulsed, or broadband). In addition, the Mel scale mimics the logarithmic perception of pitch by mammalian ears and is therefore well-suited to defined meaningful perceptual categories for mammals. Fuzzy clustering is a soft classification approach. It does not assign samples to a single category, but rather describes their position relative to overlapping categories. This method is capable of identifying stereotyped vocalisations – vocalisations located in a single category – and graded vocalisations – vocalisation which lie between categories – in a quantitative way. We evaluated the performance of this procedure on a set of long-finned pilot whale (*Globicephala melas*) calls. We compared our results with a call catalogue previously defined through audio-visual inspection of the calls by human experts. Our unsupervised classification achieved slightly lower precision than the catalogue approach: we described between two and ten fuzzy clusters compared to 11 call types in the catalogue. The fuzzy clustering did not replicate the manual classification. One-to-one correspondence between fuzzy clusters and catalogue call types were rare, however the same sets of call types were consistently grouped together within fuzzy clusters. There were also discrepancies between both classification approaches, with some catalogue call types being consistently spread over several fuzzy clusters. Compared to manual classification, the fuzzy clustering approach proved to be much less time-consuming (days vs. months) and provided additional quantitative information about the graded nature of the vocalisations. We discuss the scope of our unsupervised classifier and the need to investigate the functions of call gradation in future research.

**Author summary:** There is no consensus on how to describe the vocal repertoire of a species, an essential initial step to analyse how animals rely on different types of vocalisations according to social, ecological, and behavioural contexts. This task is even more challenging for species with graded vocal repertoires: their vocalisations do not fall into distinct categories but form a continuum which makes it difficult to draw strict boundaries between sound types. We present here a method to overcome this challenge using an unsupervised classification algorithm based on Mel frequency cepstral coefficients and fuzzy clustering. It is specifically designed to deal with the graded nature of animal vocalisations, as it can describe overlapping categories in a quantitative way. We tested our classification procedure on a particularly challenging set of long-finned pilot whale (*Globicephala melas*) calls. Indeed, this species can produce sounds of various acoustic natures (tonal, broadband, and pulsed) and their large vocal repertoire is a mix of stereotyped and highly graded sound types. We compared our results with an existing call catalogue established by human operators. We obtained promising results and recommend similar classification procedures in future studies to take a quantitative approach when studying the gradation of animals’ vocal repertoires.

## Introduction

The description of animals’ vocal repertoires can reveal vocal differences between species (1), groups (2), individuals (3), and ecological contexts (4). However, there is no consensus about the methods best suited for the construction of vocal repertoires, i.e. the features used to characterise vocalisations and the classification procedure. Various classification schemes can achieve a high degree of precision and accuracy when species produce stereotyped vocalisations (e.g. 5). However, the sounds of many species do not form discrete categories, but rather vary along a continuum, rendering categorisation challenging. Such graded repertoires appear particularly common in mammalian species (e.g. 6–8) and are also found in other taxa (e.g. birds 9).

The set of parameters used to describe the vocalisations can be a limiting factor for classification. In several studies, a reduced number of parameters in the time and frequency domains was selected (e.g. 10). This approach targets parameters that are expected to be important for categorisation and that may vary with the experimenter, the study species, or the scope of the research project. In addition, a low number of parameters may limit the detection of subtle or localized differences between vocalisations (11), which may be crucial in the study of animals’ perception of sounds. By contrast, other studies described animal vocalisations using extensive sets of parameters, such as fine-scale acoustic features (e.g. 12) or image descriptors extracted from spectrograms (e.g. 13). Such large datasets incur high computational costs. Moreover, they may require data reduction procedures such as principal component analysis (e.g. 14) or may include dimension augmentation steps (e.g. 15). The high number of parameters and their eventual transformation obscure the nature of the differences between sound classification categories and their perceived significance by study animals.

The use of Mel frequency cepstral coefficients (MFCC) as a classification parameter represents an efficient way to compress information about spectral energy distribution of a time-segment within a vocalisation with a small number of variables (16). MFCC have been widely used in speech segmentation and speech recognition, and increasingly to study animal vocalisations (e.g. 17,18). MFCC can be calculated for both harmonic and unvoiced sounds. They tend to be uncorrelated, and therefore well-suited for classification algorithms. In addition, they integrate the logarithmic scale of pitch perception by mammalian species. Categories defined on the basis of MFCC differences should thus correspond to perceptual categories, at least for mammalian species.

The procedure used to classify sounds can be another limiting factor for the description of vocal repertoires. Historically, classification has relied on the audio-visual inspection of vocalisations by trained human operators. Such methods are still commonly used (e.g. 19). They define subjective boundaries between categories based on operator inspection, which can make them difficult to precisely replicate across studies. Moreover, the operator-dependent inspection process is time-consuming and cannot easily be applied to large datasets. Supervised classification algorithms can speed up the classification. These algorithms (e.g. 2) are trained on sorted datasets, and then generalize the classification rules to new data. They allow fast and reproducible classification of animal vocalisations, although they still rely on a first-step categorisation by human operators and cannot recognize call types which are absent from their training data. Unsupervised algorithms (e.g. 20) derive classification rules from the data itself, which removes both the need for and limitations of prior human-based classification.

Most classification procedures attempt “hard” categorisation, that is to assign vocalisations to mutually exclusive categories (21). The graded nature of vocalisations, which translates into an overlap across apparent categories, is problematic for such classification approaches. Graded vocalizations increase the subjectivity and reduces the reproducibility of audio-visual categorisation by human operators. Supervised classification algorithms, which rely on geometrical rules to assign vocalisations to a category, are also sensitive to gradation. For instance, support vector machines, a commonly supervised classification algorithm (e.g. 2), compute a hyperplane in the feature space to separate categories. They aim at minimizing the number of classification errors and maximizing the distance between categories in the training dataset. When the distribution of categories overlap, a unique optimal hyperplane may not exist or may show an overly complicated shape without acoustical significance.

The fuzzy clustering (FC) algorithm used herein is a “soft” classification procedure (22). It first defines the fuzzy cluster centres as apparent stereotypes from the distribution of acoustic features in the dataset. Then, it quantifies the graded nature of vocal repertoires as the positions of individual vocalisations in-between the previously identified stereotypes. Each vocalisation is given a membership score to each fuzzy cluster. This membership score corresponds to the similarity between the vocalisation and the fuzzy cluster stereotype and can be considered as the probability that the vocalisation belongs to the fuzzy cluster. This FC algorithm has already been used successfully to classify primate and cetacean calls (21,23).

In the present study, we present a new approach for animal sound classification, based on FC and MFCC, specifically designed to deal with the graded nature of animal vocalisations. We evaluated the performance of this FC- and MFCC-based classifier on a set of long-finned pilot whale (*Globicephala melas*) calls. This dataset has previously been classified according to an expert-defined call catalogue (19), which we used for comparison. The long-finned pilot whale is a delphinid species which lives in cohesive social groups all year long (24) and has a high vocal activity in almost all behavioural contexts (25,26). Pilot whales produce a large variety of sounds including a mix of tonal, broadband, and pulsed sounds, as well as intermediate forms containing some similarities or smooth transitions from one sound type to another (19,27,28). Pilot whale calls can also be biphonated: they can contain two independently-modulated tonal components (19,29). A portion of their vocal repertoire consists of stereotyped tonal and pulsed call types. Audio-visual inspection of recordings described 125 call types, with 29 further subtypes, in long-finned pilot whales (19) and 173 call types in short-finned pilot whales *G. macrorhynchos*, a sister species (27). However, some long-finned pilot whale calls have no tonal component, and there is apparent gradation within and across call types (30–33). Long-finned pilot whales have a complex vocal repertoire, with both stereotyped and graded sound types, particularly challenging for “hard” classification. This makes them an ideal model species to test our proposed unsupervised “soft” classifier.

## Material and methods

### Animal sound dataset

We used a set of long-finned pilot whale calls recorded in northern Norway in summer 2003 using a hydrophone lowered from a small research vessel (details in 19). Calls were sampled at or downsampled to 48 kHz, and bandpass-filtered between 1000 and 22,000 Hz using a 4^th^ order Butterworth filter. Based on the audio-visual inspection of spectrograms, trained operators identified 278 high quality calls, and classified this call dataset according to a call catalogue. The calls of this dataset were assigned to eleven call types from which two were divided into 6 subtypes (Fig 1).

**Fig 1.**
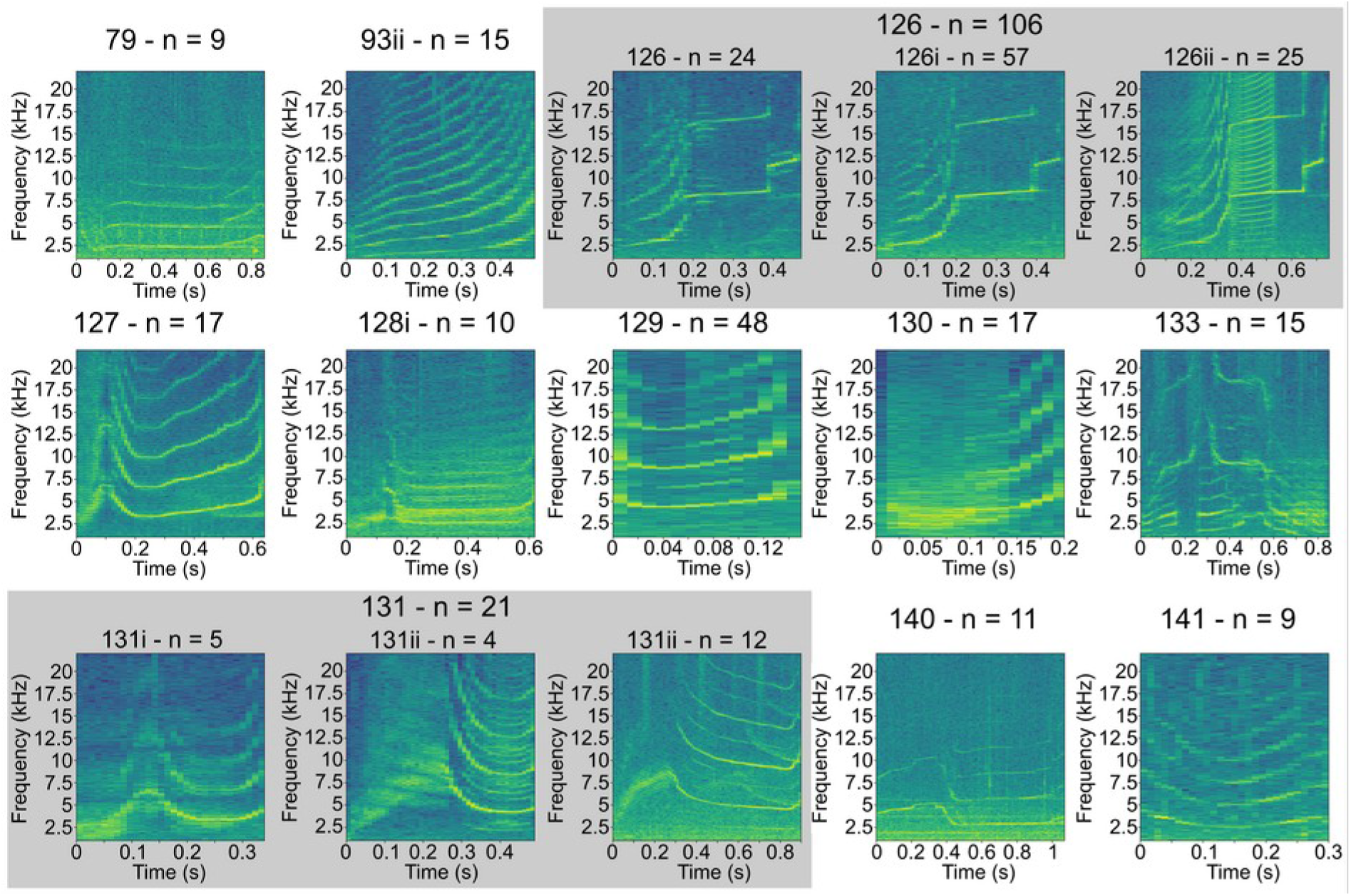
Example spectrograms of the eleven call types represented in the dataset. The spectrograms were computed with 1024 points Hanning windows and 50% overlap. The custom names of the call types are indicated as numbers and letters in large font above each spectrogram, followed by the number of samples present in the dataset (e.g. top left, n=9 calls for the call type named as “79”). Call type 126 and 131 were further divided into three subtypes, indicated by a gray background. The names and number of samples of the call subtypes are indicated in smaller font and in italics below the call type. Spectrograms and Mel energy power spectra for all vocalisations are available in the Supplementary materials File S1.

### Acoustic parameter extraction

We calculated Mel frequency cepstral coefficients (MFCC) to describe the calls (Fig 2). To do so, we created a bank of overlapping Mel scale triangular filters. Between 1000 and 22,000 Hz, we constructed 42 points linearly spaced on the Mel scale by using the Hz to Mel conversion as follows:

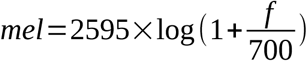

**Fig 2.**
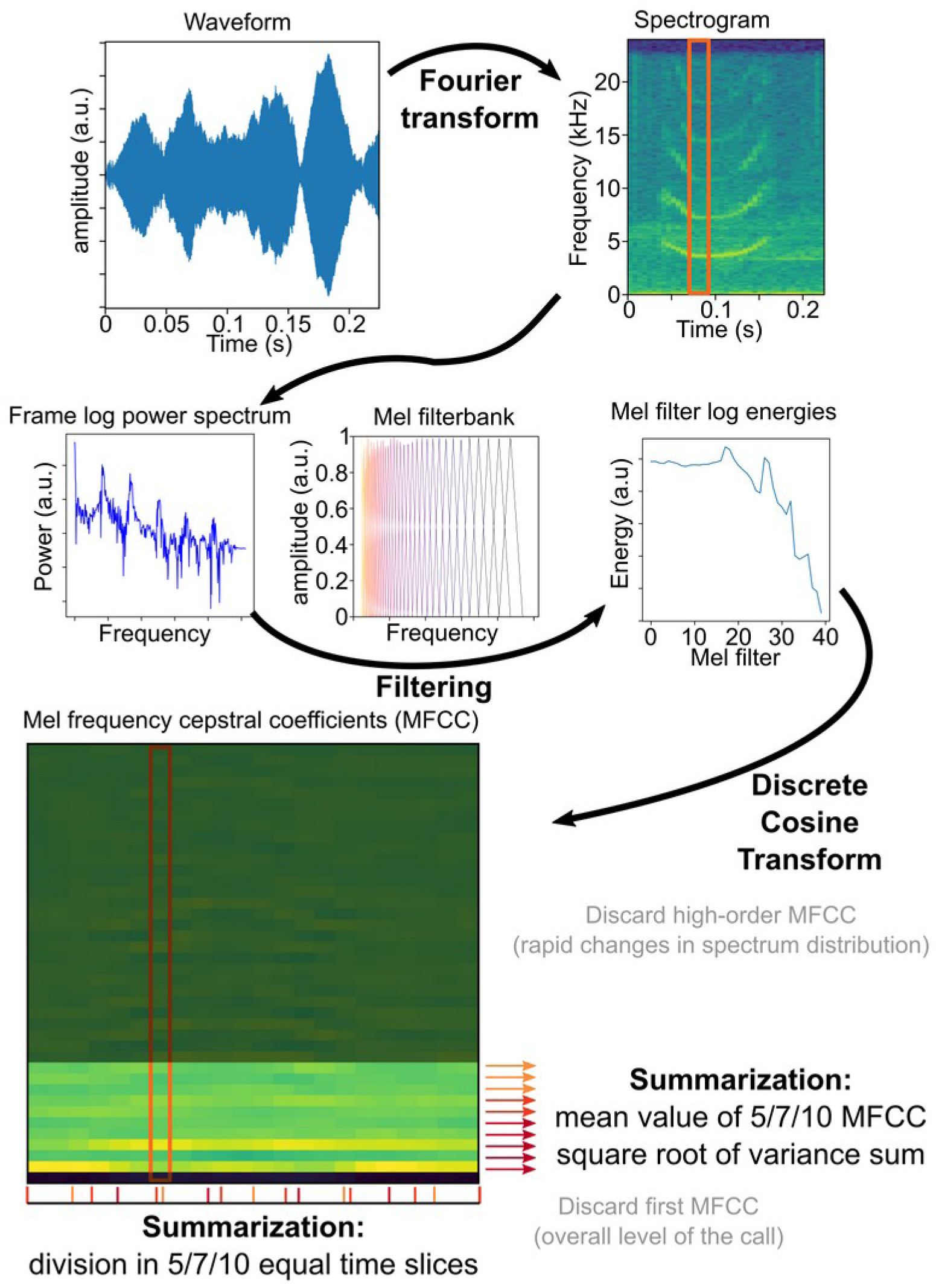
Successive steps for the calculation of the Mel frequency cepstral coefficients (MFCC). We constructed nine different datasets in the summarisation step to explore the trade-off between the descriptive power and the size of the descriptive feature set.

**mel** is the frequency in Mels and **f** is the frequency in Hz. We designed 40 overlapping triangular filters which lower and upper bounds of each filter matched the centre frequency of the adjacent frequency filters. All filters had the same weighting at their centre frequency.

We separated the calls into segments of 1024 data points, with an overlap of 50% between successive segments. We measured the distribution of energy along the frequency spectrum for each of these segments with a Fourier transform followed by a log transformation. We summed the log energy of each call segment through the 40 Mel filters, and performed a one-dimension discrete cosine transform on the 40 Mel filter log energies to obtain 40 MFCC. We discarded the first MFCC bin which corresponds to the average level of the call. We also discarded high-order MFCC bins as they correspond to rapid changes in the energy distribution along the spectrum which are less meaningful from a biological perspective.

We then summarised the MFCC (Fig 2) to use an identical number of acoustic features to describe each vocalisation. To do so, we divided each call into an equal number of time slices, and measured the mean and variance of each MFCC bin over these slices. We kept the average value of each MFCC over each time slice. We combined the variances into a single variability measure – the square root of the summed variances. The maximum amplitude of the calls was normalized before the calculation of the MFCC. MFCC values ranged roughly from minus 1 to 1. Therefore, we normalized the variability measure to the same interval using min-max normalisation.

The fuzzy clustering (FC) algorithm (further detailed below) used for call classification is sensitive to high dimensionality (34,35). When using a large number of descriptive features, FC tends to assign all samples to a single fuzzy cluster, located around the centre of gravity of the dataset. We built nine datasets with different numbers of time slices per call to investigate the trade-off between the descriptive power of the features set and the ability of the FC to detect multiple categories. We kept either five, seven, or ten MFCC bins, summarised over five, seven, or ten time slices. The number of features in these sets ranged from 26 to 101. We named each set of features as **“XMYS”** with **X** being the number of MFCC (**“M”**) and **Y** corresponding to the number of time slices (**“S”**). As an example, **“5M7S”** corresponds to the features set with five MFCC summarised over seven time slices.

We used the Python package “pylotwhale” (https://github.com/floreencia/pylotwhale) to extract the MFCC measurements from the pilot whales’ calls.

### Unsupervised classification procedure

The FC algorithm is an unsupervised classifier, which here defines fuzzy clusters from the distribution of acoustic features (MFCC) extracted from the dataset. The algorithm detects stereotypes in the dataset, around and between which individual calls appear to be distributed. These apparent stereotypes represent the centre of the fuzzy clusters, which are then used to describe a “soft” partition of the dataset.

Rather than assigning a given call to a single category, the FC algorithm provides for each call a membership score to each fuzzy cluster. These membership scores reflect the similarity between the call and the centre of the fuzzy clusters. Membership scores range from 0 to 1, and the sum of all the membership scores of a call is equal to one. Therefore, membership scores are akin to the probability that a call belongs to a given fuzzy cluster and allow for the quantification of the graded nature of the vocal repertoire. For instance, assume that a hypothetical vocal repertoire consists in fuzzy clusters A and B. A very stereotypical vocalisation may have a membership score of 0,95 for A and 0,05 for B, meaning that it is very similar to the stereotyped template of A. By contrast, a graded vocalisation may have a membership score of 0,35 for A and 0,65 for B, meaning that it lies between the stereotyped templates of A and B but is somewhat closer to the template of B than the template of A.

There exist a multitude of fuzzy partitions for a given dataset. These partitions differ in the number of categories (noted as **k**) and the fuzziness, i.e. the amount of overlap between neighbouring categories (noted as **p**). **p** and **k** are the two key parameters which drive the FC algorithm (21,22,36). The fuzziness quantifies the extent to which fuzzy clusters are allowed to overlap: a fuzziness of 1 corresponds to no overlap between clusters (equivalent to “hard” clustering). The higher the fuzziness value, the more overlap there can be between clusters.

For a given **(p, k)** pair, the FC algorithm performs the following four steps:

1. Initialize **k** random fuzzy cluster centres. We drew the coordinates of the initial cluster centres from uniform distributions over the range of each feature in the call dataset.
2. Iteratively calculate membership scores from the distances between data points and cluster centres, taking the fuzziness value **p** into account (see equation 1), and update the positions of the cluster centres based on membership scores, data points, and fuzziness (see equation 2).
3. Repeat step 2 until obtaining the convergence of an objective function (equation 3). The objective function, namely the sum of the squared distances between data points and cluster centres weighted by membership scores and fuzziness, is also a measure of cluster compactness. We used it as a measure of clustering quality. We stopped the iterative process when the objective function decreased by less than 0.0001.
4. Check the distances between fuzzy cluster centres. If some fuzzy cluster centres differed by less than 1% of the range of every feature, we fused them together. The centre of the fused cluster was defined as the mean of the fused clusters’ centres. We updated membership scores (equation 2) after fuzzy cluster fusion.

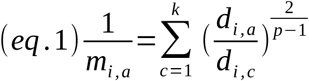

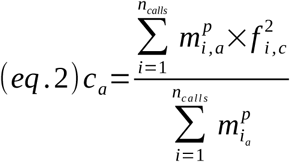

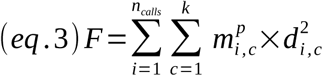

With:

– **m_i,a_** the membership score of vocalisation **i** to fuzzy cluster **a**
– **k** the number of fuzzy clusters
– **d_i,a_** the Euclidian distance between vocalisation **i** and the centre of fuzzy cluster **a**
– **p** the fuzziness value
– **c_a_** the coordinates of the centre of fuzzy cluster **a** in the feature set space
– **f_i_** the coordinates of vocalisation **i** in the feature set space
– **F** the value of the objective function

The FC algorithm can be sensitive to local minima of the objective function. To avoid the detection of local minima instead of the global minimum, we ran the algorithm 100 times for each **(p, k)** pair and kept the outcome with the lowest final value of the objective function.

The full classification procedure started with the selection of the maximal number of clusters allowed **k_max_** and the initial value of fuzziness **p_start_**. **k_max_** should be larger than the number of expected clusters in the dataset so as not to be a limiting factor for classification. **p_start_** should be chosen so that the FC results in a single fuzzy cluster. Both **k_max_** and **p_start_** should not be too large to avoid additional computational costs. Taking these constraints into account, we set up **k_max_** = 15, as there were 11 catalogue call types in the dataset, and **p_start_** = 2.5 based on pilot tests. We then ran the FC algorithm for **p = p_start_** and all the values of **k** between 2 and **k_max_**, and we kept the clustering solution with the lowest objective function. We then decremented **p** by 0.01 and ran the algorithm again until **p** = 1.01.

### Fuzzy clustering analyses

#### Identification of valid clustering solutions

The FC procedure produces one partition of the dataset into fuzzy categories for each **(p, k)** pair. Therefore, the first step of the analysis of FC results is the identification of interesting clustering solutions.

For high values of fuzziness, the algorithm describes a single fuzzy cluster containing the entire dataset. As fuzziness decreases, the fuzzy clusters get separated and their degree of overlap diminishes. Clustering solutions which are optimal over large ranges of fuzziness are likely to represent underlying structures in the dataset.

For each of the nine features sets, we plotted the optimal number of clusters as a function of the fuzziness value to visually identify the optimal clustering solutions. We then verified that these clustering solutions were consistent. That is, we checked that the partitions of the dataset having the same number of fuzzy clusters but obtained with different values of fuzziness did not result in markedly different clusters.

To visualize the clustering solutions, we used PCA (principal component analysis) and UMAP (uniform manifold approximation and projection). PCA is a standard procedure for dimension reduction and data visualisation. We drew 70% confidence ellipses to visualise either the repartition of the fuzzy clusters or the repartition of the catalogue call types over the dataset on the PCA plots. However, the distance between data points on the PCA projection does not faithfully reflect their distance, because the main objective of PCA is to conserve the variance of the dataset. UMAP is a data reduction procedure which finds a low-dimension, non-linear projection of the dataset which is as close as possible to the high-dimension topology of the dataset (see 37 for an application in bioacoustics). This means that the distance between samples on the UMAP projection should be more similar to the Euclidian distance between these points than on the PCA projection. PCA and UMAP projection plots, coloured by fuzzy cluster or by catalogue call type, can be found in the Supplementary materials Files S2 and S3.

#### Harmonisation of the FC solutions

Since the initialisation of the fuzzy cluster centres was random, the order of the fuzzy clusters was not necessarily consistent across the different realisations of the algorithm. Similarly, the order of fuzzy clusters in solutions having the same number of clusters but obtained using different feature sets may also be different. The harmonisation of fuzzy cluster order was thus required before further analyses.

To harmonise fuzzy cluster order across fuzziness values, we used the lowest value of fuzziness as a reference. The partition for this reference value of fuzziness allowed the lowest amount of overlap between fuzzy clusters and should therefore exhibit the clearest separation of fuzzy clusters. We then compared the main fuzzy cluster i.e. including the call with the highest membership score, among all other FC realisations having this reference, and we adjusted the order of the fuzzy clusters to maximize similarity.

We used a similar strategy to assess the best correspondence between fuzzy clusters obtained from different feature sets. We verified our harmonisation procedure visually by assessing the relative positions of fuzzy clusters on PCA and UMAP plots (Supplementary materials File S2 and S3). We removed the FC realisations which were markedly different from the other ones in the clustering solutions.

#### Characterisation of the fuzzy clusters

After the selection of optimum clustering solutions, we aimed to characterize the centre of the fuzzy clusters which correspond to the stereotypes around which the vocal repertoire is organised. We reconstructed a stereotyped template for each fuzzy cluster, which reflects the hypothetical call corresponding to the centre of the fuzzy cluster.

In order to do so, we inverted the MFCC computation procedure (Fig 2). Starting from the coordinates of the fuzzy cluster centre in the feature set space:

– We discarded the variability measure.
– We zero-padded the MFCC for each time slice to replace the discarded MFCC.
– We performed an inverse discrete cosine transform on the MFCC, which provided an approximation of the Mel filter log energies.

This procedure builds an approximation of the hypothetical call spectrograms corresponding to the centre of the fuzzy clusters (see 38 for a similar approach), which should represent the stereotyped calls around which the vocal repertoire is distributed. This method, far from a full audio reconstruction pipeline, warranted caution in the interpretation of the reconstructed calls. First, the spectral and temporal resolution of the reconstructed vocalisations depended on the feature set used. The reconstructed vocalisations were based on the same time slices as the MFCC feature sets and the higher the number of MFCC, the finer the spectral resolution of the reconstructed vocalisations. Second, we discarded the first MFCC which encoded the average sound pressure level of each time slice. The reconstruction provided an overall distribution of energy along the frequency spectrum of the call, but no indication about potential variations of amplitude.

#### Gradation analysis

The reconstruction of the stereotypical calls characterizes the centre of the fuzzy clusters, but gives no indication regarding the distribution of calls around these stereotypes. To study the gradation of the vocal repertoire of long-finned pilot whales, we measured the typicality of their calls which corresponds to the difference between the two highest membership scores of a given call. A stereotyped call will be much closer to the template of its main fuzzy cluster (i.e. the fuzzy cluster to which it has the highest membership score) than to the templates of other fuzzy clusters, resulting in a high typicality value. By contrast, a graded call will have similar membership scores to at least two fuzzy clusters resulting in a low typicality value.

In order to visualize call gradation between fuzzy clusters, we plotted the membership score to a fuzzy cluster relative to the membership score of another fuzzy cluster (as in 21). The position of individual calls in these triangular plots reflects the gradation between the two fuzzy clusters in the dataset (Fig 3). For each pair of fuzzy clusters, we decided to only represent the calls for which the two highest membership scores corresponded to these two fuzzy clusters; this reduced the number of calls displayed on the plot and eased the readability of the figures. As an example, if the two highest membership scores of a call were associated to clusters A and B, then this call would be only represented on the triangular plot of clusters A and B, but not on the triangular plot of any other pairs of clusters such as clusters A and C. Triangular plots of all analysed calls can be found in the Supplementary materials File S4.

**Fig 3.**
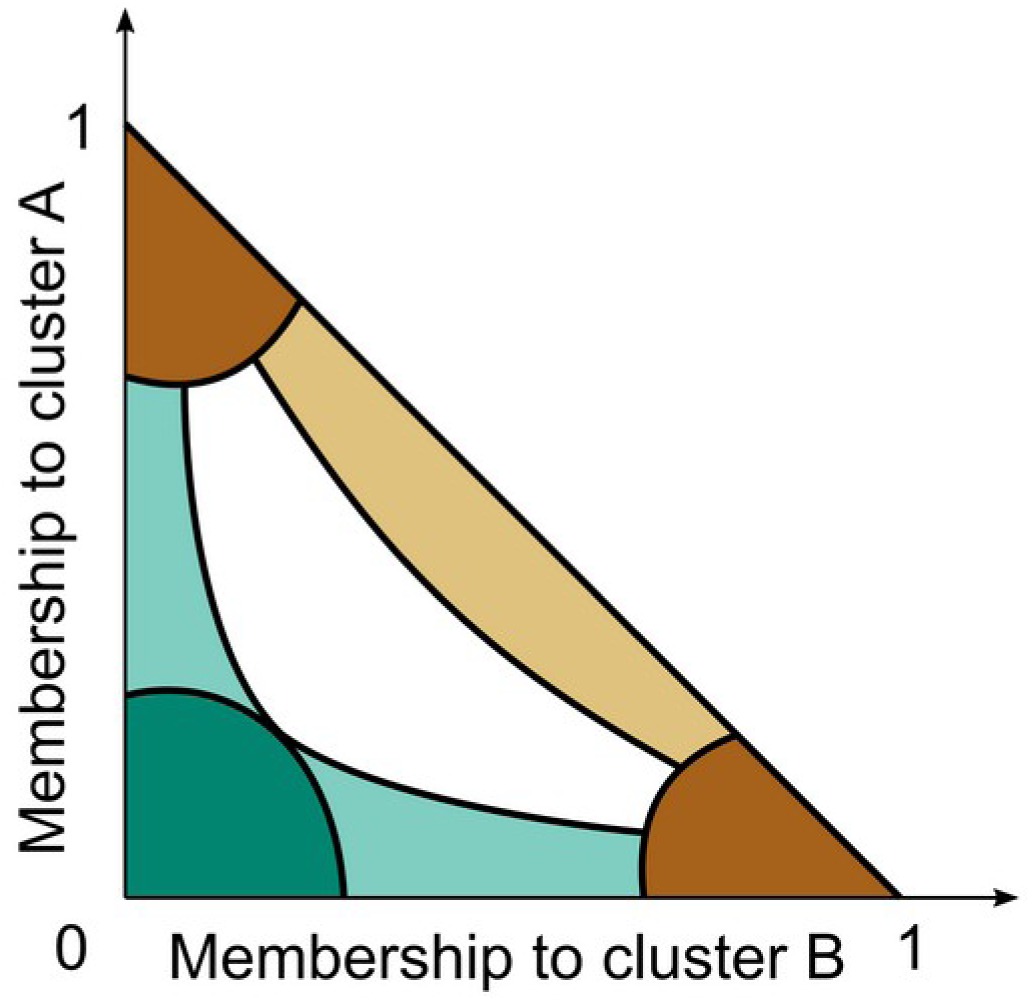
Triangular gradation plots providing a detailed analysis of the call gradation within a pair of clusters. Highly stereotyped calls, in dark brown, are around the tips of the triangular plot: they have a very high membership to one cluster, and low memberships to all others. Calls along the diagonal of the triangular plot, in light brown, are graded almost exclusively between clusters A and B: they have varying memberships to these clusters, and the sum of their membership scores to A and B is close to 1. Calls located along the vertical (or horizontal) side of the triangular plot, in light green, are graded between cluster A (or B) and a third cluster: they have varying membership to cluster A (or B) and consistently low membership to cluster B (or A). Calls close to the origin of the triangular plot, in dark green, mostly resemble other clusters: they have low memberships to both A and B. Calls located in the central area of the triangular plot, in white, are graded between several clusters, including A and B.

#### Comparison between the FC and the catalogue-based classification

We investigated the performance of our FC procedure by comparing the obtained fuzzy clusters with the call types described previously in a call catalogue established by a human operator. We constructed contingency tables crossing the outcomes of both classifications. We faced two challenges in doing so. First, we had to decide how to translate membership scores into counts in the contingency tables, since the calls were not assigned to a single fuzzy cluster. Second, the FC solutions were not unique and clearly defined. Each clustering solution was obtained i) for multiple values of fuzziness and ii) with one or several sets of descriptive features. To overcome these challenges, we calculated average membership scores over all the realisations of each FC solution and used them as counts in the contingency tables. Therefore, the higher and more consistent the membership scores of a call to a fuzzy cluster, the higher the corresponding value in the contingency table. We used the PCA and UMAP plots (Supplementary Materials File S2 ans S3) to visually compare the outcome of the FC procedure and the catalogue-based, human-operated classification.

We developed a Python package “fuzzyClustering” to run the FC algorithm and analyse the clustering results (code available at https://github.com/benbenti/fuzzyclustering). Module “algorithms” runs the FC classification and measures typicality, and module “visualisation” contains all the visualisation tools used to produce the figures in the Results section. We used the modules “umap-learn”, “preprocessing” and “decomposition” of Python package “sickit-learn” (37) for the UMAP and PCA.

## Results

### Identification of stable FC solutions

The FC algorithm that we developed was able to reach convergence for all **(p, k)** pairs and for all feature sets. The number of loops required to reach convergence was similar for all feature sets. It ranged from 4 to 184 and had an average of 73 +/− 31 loops (mean +/− SD). As expected, for low values of fuzziness, the algorithm resulted in clustering solutions with the maximal number of fuzzy clusters (**k_max_** = 15) using all feature sets (Fig 4). As fuzziness increased, the number of fuzzy clusters in the optimal solution decreased, reaching one for the largest feature sets (“**10M”** feature sets, Fig 4). Indeed, we identified stable clustering solutions for all feature sets (Fig 4).

**Fig 4.**
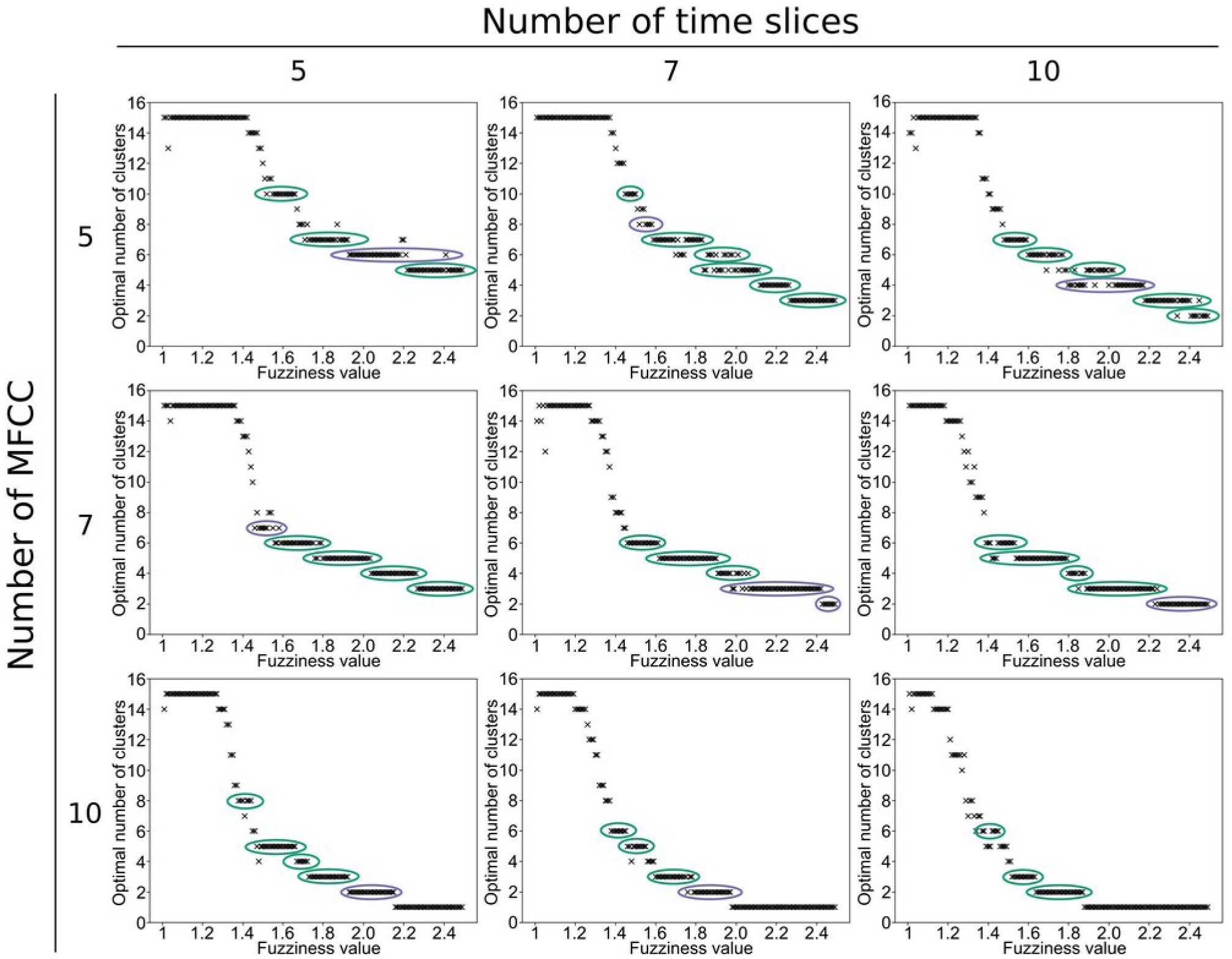
Optimal number of clusters relative to fuzziness for all sets of descriptive acoustic features of the calls. Stable clustering solutions are circled in green when all the realisations (i.e. the clustering solutions for the different fuzziness values) were strongly consistent and in purple when there were slight variations between the realisations, as evaluated by visual inspection of the PCA and UMAP plots (Supplementary materials Files S2 and S3). Discarded realisations are not included in the coloured circles.

Some dataset partitions were optimal over large intervals of fuzziness (Fig 4, Table 1). Visual inspection of PCA and UMAP plots (Supplementary materials Files S2 and S3) showed that the different realisations of the FC algorithm within these fuzziness intervals, that is optimal clustering solutions with the same number of fuzzy clusters but different values of fuzziness, were highly consistent.

**Table 1.**
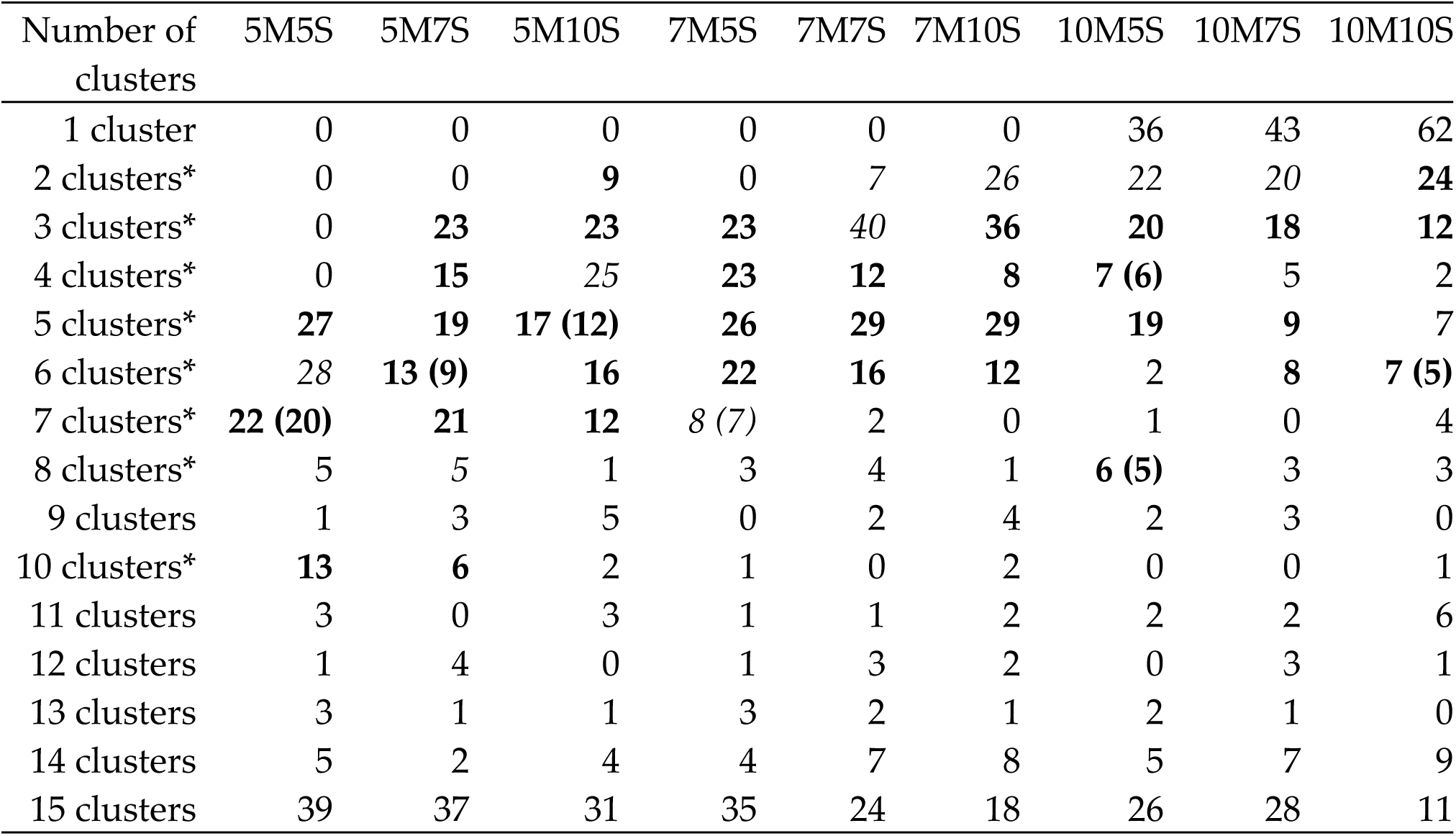
Number of fuzziness values for which the different FC solutions were optimal. Bold cells correspond to FC solutions which are strongly consistent across fuzziness values (green circles in Fig 4). Cells in italics correspond to FC solutions which showed slight variations across fuzziness values (purple circles in Fig 4). Numbers in brackets correspond to the number of realisations selected for further analysis after the removal of inconsistent realisations. *: clustering solutions selected for further analyses.

A few realisations (15 out of 768) described dataset partitions markedly different from realisations with close fuzziness values. We discarded these realisations from further analyses. The discarded FC realisations were: **“5M5S”** – p=2.19 and p=2.20; **“5M7S”** – p=1.70, p=1.72, p=1.73, and p=1.74; **“5M10S”** – p=1.48, p=1.69, p=1.75, p=1.78, and p=1.79; **“10M5S”** – p=1.38 and p=1.48; **“10M10S”** – p=1.34 and p=1.37.

The FC procedure produced similar results with the nine feature sets. Taking into account both the number of realisations and their consistency, we selected clustering solutions having two to six fuzzy clusters for further analyses (Table 1). These clustering solutions were detected using all feature sets except the largest and the smallest ones (Fig 4). This stability reflected prominent structures in the dataset (Fig 4, Table 1). We also analysed clustering solutions with seven, eight, and ten fuzzy clusters. These clustering solutions were detected using fewer feature sets and showed less consistency across fuzziness values.

### Characterisation of the fuzzy clusters

For each feature set, clustering solutions with the same number of fuzzy clusters but different values of fuzziness led to the reconstruction of highly similar stereotypes (Supplementary Materials Fig S5 and S6). This result confirmed that the stable solutions of the FC procedure described the same fuzzy clusters over a range of fuzziness values. We were able to identify, from the reconstructed stereotypes, which solutions were highly consistent and which showed slight variations across realisations (Fig 4). All reconstructed vocalisations can be found in the Supplementary materials File S7.

The overall shape of the reconstructed cluster centres was similar when using different feature sets, despite the differences in spectral and temporal resolutions (Supplementary Materials Figs S5 and S6, File S7). This means that the FC procedure detected the same structures in the dataset using descriptive feature sets of different temporal and spectral precisions.

Stereotypes reconstructed from **“5M”** feature sets showed the lowest resolution: we could only describe the dataset as a continuum from low frequency clusters – cluster centres with most energy in a frequency band centred around the 11^th^ and 12^th^ Mel filter (800-1300 Hz) – and mid-frequency clusters – cluster centres with most energy in a frequency band centred on the 25^th^ Mel filter (5-6 kHz). All **“5M”** reconstructed stereotypes presented both frequency bands with varying relative amplitude (Supplementary Materials Fig S5).

Increasing numbers of MFCC and time slices resulted in a more precise reconstruction of the cluster centres. The **“10M10S”** six-cluster solution (Supplementary Materials Fig S6) was the solution with both the highest number of fuzzy clusters and the highest precision in the call reconstruction. This reconstruction revealed the same continuum between low frequency and high frequency stereotypes we observed with the **“5M”** feature sets. On one end of the continuum, we observed cluster centres characterised by a high energy low frequency band (centred on the fifth and sixth Mel filters, between 300-600 Hz) and a fainter element, located around the 12^th^ Mel filter (between 1,1 and 1,4 kHz) and presenting multiple harmonics. On the other end of the continuum, we observed cluster centres characterised by a slightly u-shaped, mid-frequency element (ranging from the 23^rd^ – around 4-5 kHz – to the 25^th^ cluster – around 5-6 kHz), with one harmonic band (around the 32^nd^-33^rd^ Mel filter, 10-13 kHz). These precise reconstructions allowed a better description of the intermediate cluster types.

The reconstructed stereotypes were also similar across the different stable solutions obtained with the same feature set (Supplementary Materials File S7 and Fig S8). This similarity meant that the different clustering solutions described the same structure in the dataset at different levels of precision. Visual inspection of the UMAP plots (Supplementary materials File S3) confirmed that the increase in the number of fuzzy clusters corresponded to the subdivision of previously identified fuzzy clusters. For instance, the four-cluster solution obtained with the **“7M10S”** feature set corresponded to the three-cluster solution, with one fuzzy cluster split in two (Fig 5).

**Fig 5.**
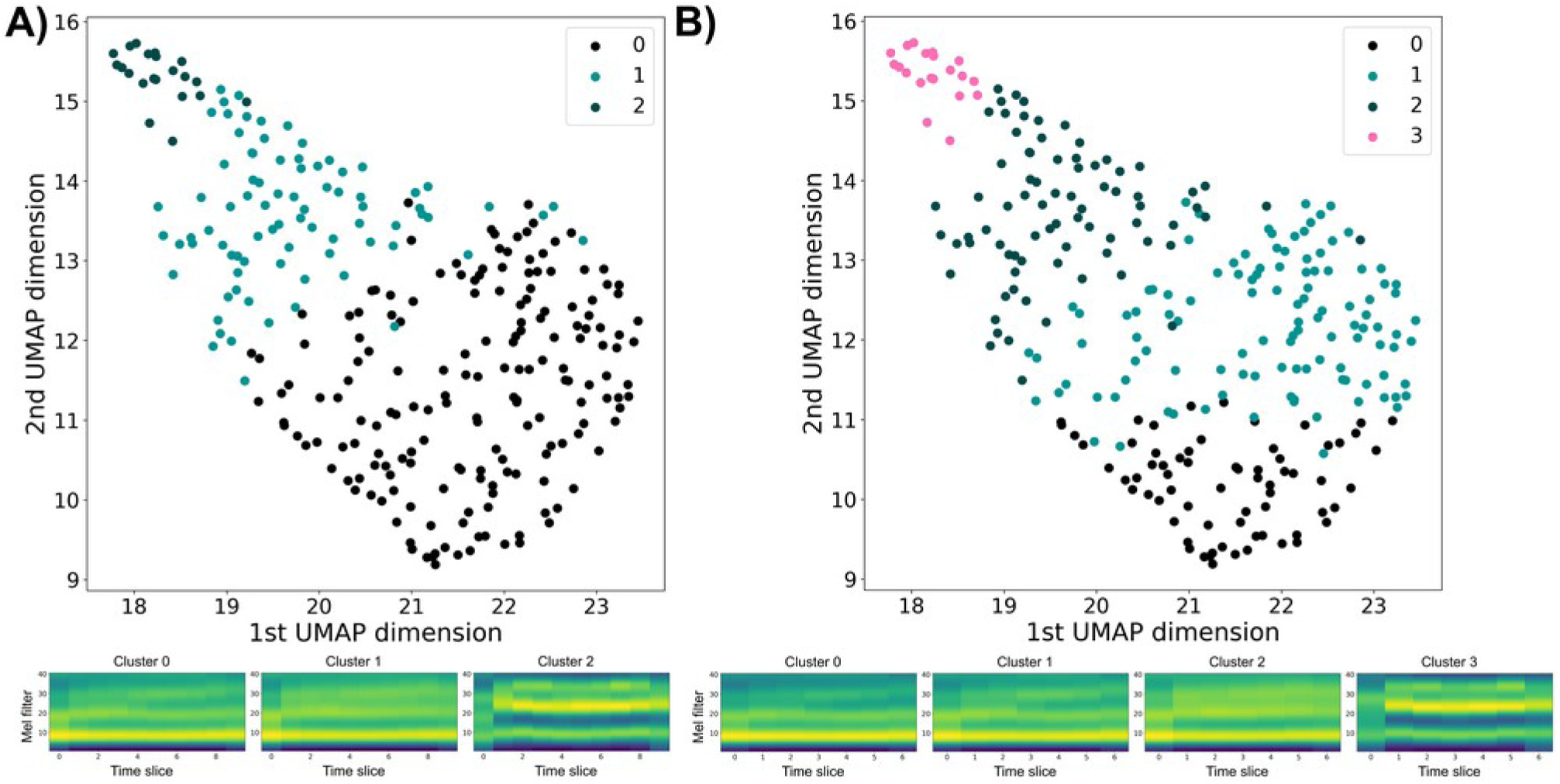
UMAP plots and reconstructed cluster centres obtained with the “7M10S” feature set. A) Three-cluster solution (p = 2,24). B) Four-cluster solution (p = 1,80). Cluster 1 and 2 in the three-cluster solution are almost identical to Cluster 2 and 3 in the four-cluster solution. Cluster 0 in the three-cluster solution is divided into Cluster 0 and 1 in the four-cluster solution.

Most clustering solutions described the dataset as a linear-like continuum from low frequency, rising frequency fuzzy clusters with many harmonic bands to mid-frequency, u-shaped fuzzy clusters with a single harmonic band. There were a few exceptions to this observation: the six-cluster solution obtained with the **“10M5S”** and **“10M7S”** feature sets, the eight-cluster solution obtained with the **“10M5S”** feature set (Supplementary Materials Fig S8), and the ten-cluster solutions. These exceptions were the clustering solutions having the highest number of clusters for each feature set.

### Gradation analyses

#### Typicality

We observed three different patterns of typicality distribution (Fig 6):

a. Clusters for which associated calls (i.e. when the highest membership score was to this cluster) had mostly low (<0.3) values of typicality.
b. Clusters for which associated calls were roughly uniformly distributed over a large range of typicality. There were calls with high typicality (>0.6) and calls with low typicality (<0.3) associated with these clusters.
c. Clusters for which associated calls showed low and intermediate values of typicality.

**Fig 6.**
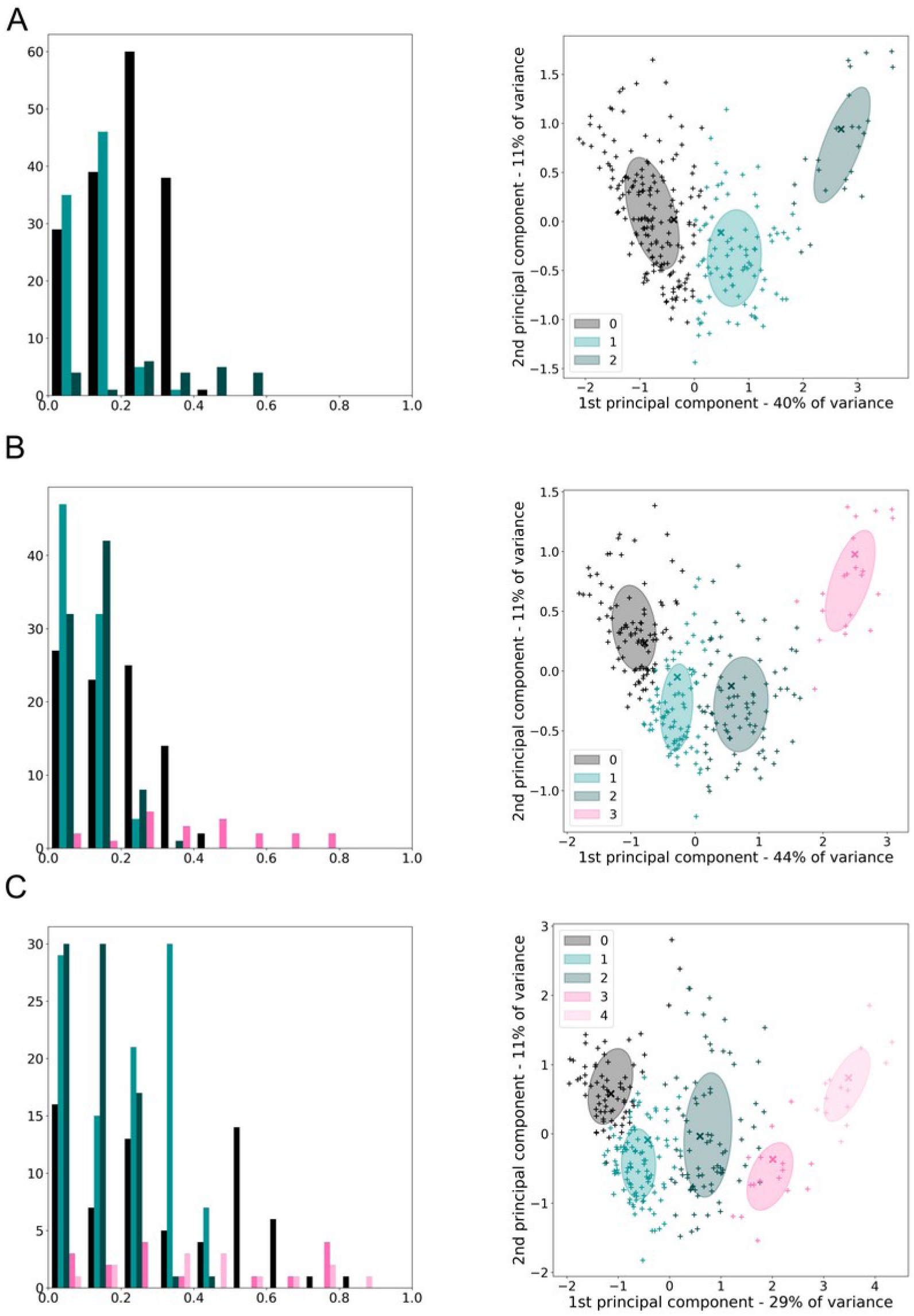
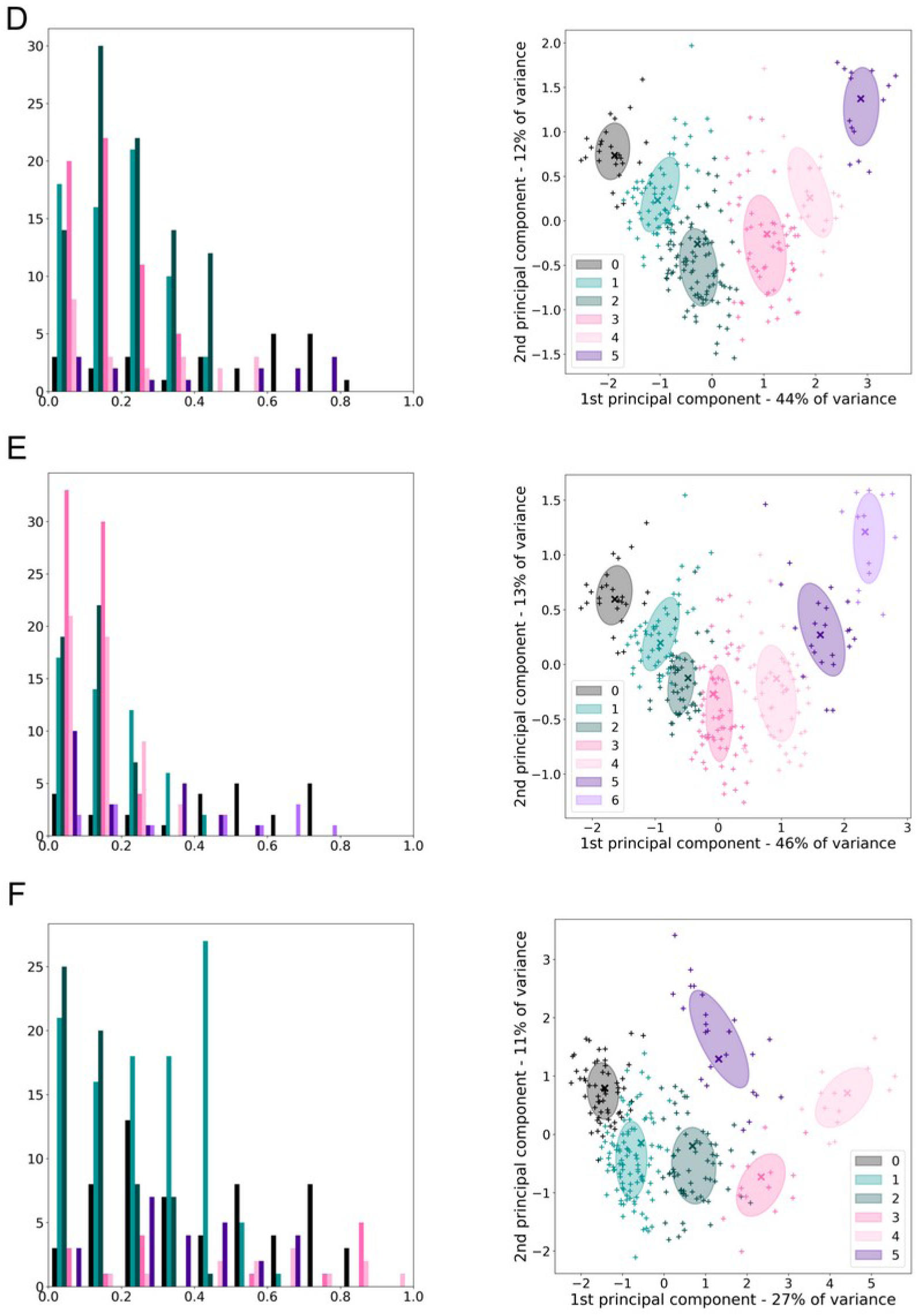
Histogram of typicality values and associated PCA plots for clustering solutions with three to eight fuzzy clusters. The left panel shows a histogram of typicality values. Each call is coloured according to its most similar fuzzy cluster, referred to as its main cluster. Calls with high typicality values are much closer to their main cluster than to the other ones (they are stereotyped). Vocalisations with low typicality values are not much closer to their main cluster than to other ones (they are graded or variable).

The right panel shows the PCA plot of the corresponding FC realisation for the interpretation of typicality values. A single realisation of the FC, chosen randomly, is presented for each clustering solution. The clustering solution with two fuzzy clusters is not represented in this figure, since typicality is devoid of sense when considering only two fuzzy clusters. Similar figures for all clustering realisations are presented in the Supplementary materials File S7. **A) “7M7S”, p = 2.27 “7M5S”, p = 2.17 C) “10M7S”, p = 1.53 D) “5M10S”, p = 1.66 E) “5M7S”, p = 1.69 F) “10M5S”, p = 1.43**

Fuzzy clusters (a) corresponded to fuzzy clusters surrounded by close neighbours. Therefore, all calls which were close to the centre of these clusters were not too far from at least one of their neighbours, resulting in low typicality values. There were no strongly stereotyped calls resembling the centre of these fuzzy clusters: they do not correspond to strong stereotypes in the call repertoire.

Fuzzy clusters (b) were located at the edge of the dataset. Calls mainly associated with these clusters were either between this cluster and their nearest neighbour, resulting in low and intermediate typicality values, or towards the edge of the dataset and therefore much closer to this cluster than to other ones, resulting in high typicality values. There were strongly stereotyped calls resembling the centre of these fuzzy clusters.

Fuzzy clusters (c) corresponded to clusters surrounded by distant neighbours. Calls associated with these clusters did not present high typicality values, but showed higher typicality values than the calls associated with clusters (a).

The density of the dataset around the centres of the fuzzy clusters influenced the distribution of typicality. Clusters which centre was located in dense areas of the dataset had more calls close to their centre: the calls mainly associated with these clusters showed higher typicality values. We observed the opposite for clusters which centre was located in sparse areas of the dataset: their centre was more distant from neighbouring calls resulting in lower values of typicality.

As fuzziness increased, the observed patterns of typicality distribution remained the same, but typicality values tended to decrease overall. Higher values of fuzziness led to more overlap between neighbouring fuzzy clusters, resulting in less clear separation of fuzzy clusters and a reduced typicality of the calls.

The range of typicality values extended upward as the number of clusters increased. Even for larger number of fuzzy clusters, calls were still located between fuzzy clusters and obtained low values of typicality. However, calls which were very close to the centre of a fuzzy cluster had their membership scores distributed over a larger number of distant clusters, which resulted in increased typicality values. In addition, clustering solutions with larger number of clusters were stable at lower values of fuzziness, which were associated with an increase in typicality.

#### Pairwise gradation

The analysis of the triangular gradation plots provided additional information about the gradation across fuzzy clusters. In the example shown in Fig 7, no call had its highest membership scores to Cluster 0 and 2, and calls were located close to the diagonal line of the triangular plots (Fig 7A), which indicates gradation between pairs of clusters with low membership to the third fuzzy cluster. Membership scores to Clusters 0 and 2 reached much higher values than membership scores to Cluster 1. These observations described the dataset as a linear continuum from Cluster 0 to Cluster 2, with Cluster 1 in the middle. The reconstructed cluster centres were consistent with this observation, with the centre of Cluster 1 being an intermediate form between the centres of Cluster 0 and 2 (Fig 7B). Some calls were very similar to the ends of the continuum (see rightmost call in Fig 7C and the reconstruction of Cluster 2 in Fig 7B), whereas the calls closest to the intermediate cluster showed low values of typicality (Fig 7C). Triangular plots for all clustering solutions can be found in the Supplementary materials File S4.

**Fig 7.**
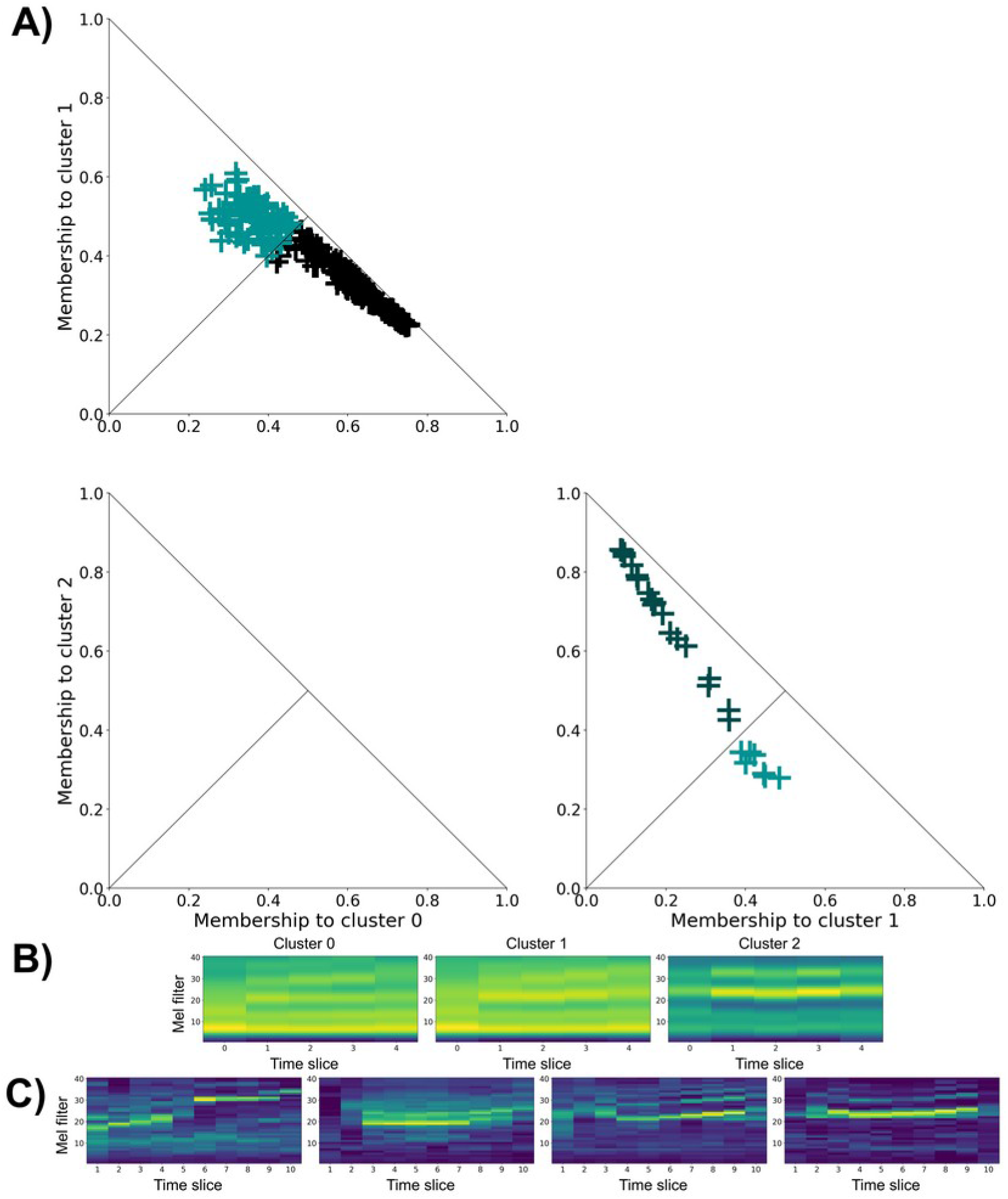
Triangular gradation plots (A), reconstructed cluster centres (B), and example spectrograms (C) for the three-cluster solution obtained with the “10M5S” feature set (p = 1,80) **A)** Each call is only represented once, on the triangular gradation plot showing the two clusters to which it has the highest memberships **C)** From left to right: the call graded between Cluster 0 and 1 closest to Cluster 0 (membership score to Cluster 0: 0.75; typicality 0.53); the call graded between Cluster 0 and 1 closest to Cluster 1 (membership score to Cluster 1: 0.52, typicality: 0.15); call graded between Cluster 1 and 2 closest to Cluster 1 (membership score to Cluster 1: 0.49; typicality: 0.21), and call graded between Cluster 1 and 2 closest to Cluster 2 (membership score to Cluster 2: 0.86; typicality: 0.77).

In most cases, calls were only present in the rightmost plot of each line in the compound figures (see Fig 7-8 and Supplementary materials File S4), indicating that the centres of the fuzzy clusters were aligned in numerical order in the feature set space. If a given call had its higher membership score to one fuzzy cluster, its second highest membership score could only be either the previous or the following cluster in the alignment (except for fuzzy clusters located at the ends of the alignment, which had a single neighbour).

**Fig 8:**
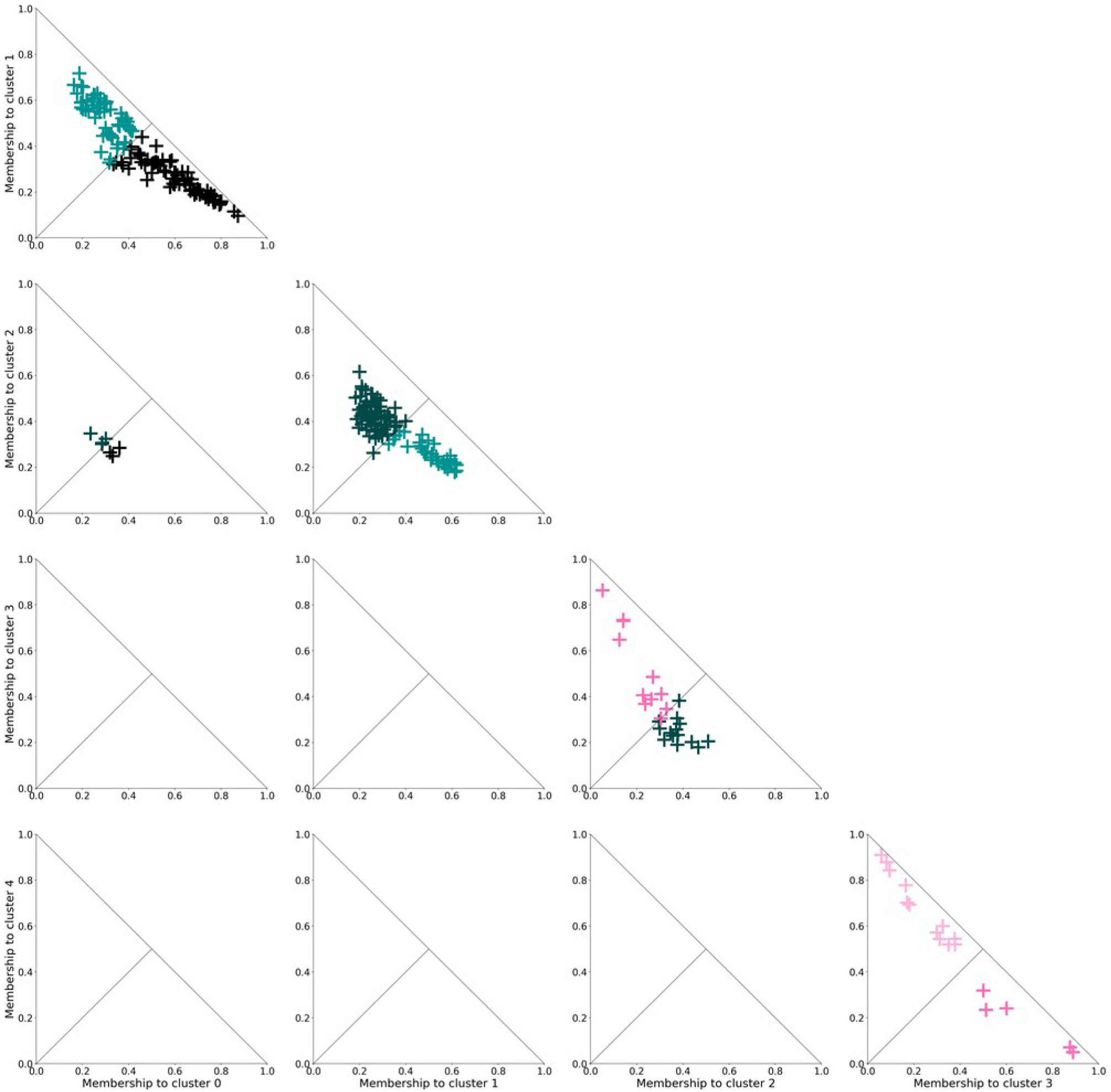
Triangular gradation plots for the five-cluster solutions obtained with the “10M5S” feature set (only one realisation shown: p = 1,56). Almost all calls (272/278) are in the rightmost plot of each line, indicating that fuzzy clusters are aligned in the numerical order in the feature set space. Membership scores reached higher values for Cluster 0, 4, and 3 than for Cluster 1 and 2.

The highest membership scores and the clearest pairwise gradation (calls located closer to the diagonal of the triangular plots in Fig 7-8) involved the fuzzy clusters located at the ends of the continuum described by the FC results. The calls were closer to the central part of the triangular plots for intermediate clusters (Fig 7-8, Supplementary materials File S4), indicating a more graded nature for these calls.

As expected, while fuzziness increased, all calls were closer to the centre of the triangular plots (see Supplementary materials File S4), indicating that calls became graded between multiple clusters for high values of fuzziness.

The number of calls displayed in the triangular plots varied across panels (Fig 8). The triangular plots showing the gradation between low-frequency fuzzy clusters (leftmost clusters in Supplementary Materials Fig S6 and S8; Cluster 0, 1, and 2 in Fig 11) contained a large number of calls, whereas the plots showing the gradation between mid-frequency u-shaped clusters (rightmost clusters in Supplementary Materials Fig S6 and S8; Cluster 3 and 4 in Fig 11) displayed few calls. In addition, the maximum membership scores were lower for the former, with the exception of Cluster 0, than for the latter (Fig 8). The dataset was denser around the centres of fuzzy clusters characterised by much energy in the low frequencies than around the centre of fuzzy clusters characterised by much energy in the mid-frequency band (see PCA plots in the Supplementary materials File S2).

There were some exception to this general pattern, with some compound figures showing calls not only in the rightmost plot of each line: the six-cluster solutions obtained with the “**10M5S”** and “**10M7S”** feature sets, the eight-cluster solution obtained with the “**10M5S”** feature set, and the ten-cluster solutions (Supplementary materials File S2). In these clustering solutions, one or several clusters seemingly lay outside of the linear alignment of fuzzy clusters and showed gradation with several neighbouring clusters.

### Comparison of fuzzy clusters with the call catalogue

We built correspondence tables for all realisations of the FC algorithm (Supplementary materials File S9). Correspondence tables were similar across fuzziness values except the two-cluster solutions obtained with the “**7M10S”**, “**10M5S”**, and “**10M7S”** feature sets (Supplementary materials File S9) for which the boundary between fuzzy clusters changed with fuzziness. The relationship between fuzzy clusters and catalogue call types wasn’t clear and varied slightly depending on which feature set had been used in the FC procedure (see correspondence tables in the Supplementary materials File S9).

The FC procedure did not replicate the manual classification of pilot whale calls. It achieved slightly lower precision than the call catalogue: we defined between two and ten fuzzy clusters, with most support for the classifications into five and six fuzzy clusters compared to 11 call types in the catalogue. Nevertheless, we observed similarities between the fuzzy clusters and the catalogue call types. An example correspondence table, associated with membership scores of individual calls and UMAP plots for visual comparison of fuzzy clusters and call types is presented in Fig 9.

**Fig 9.**
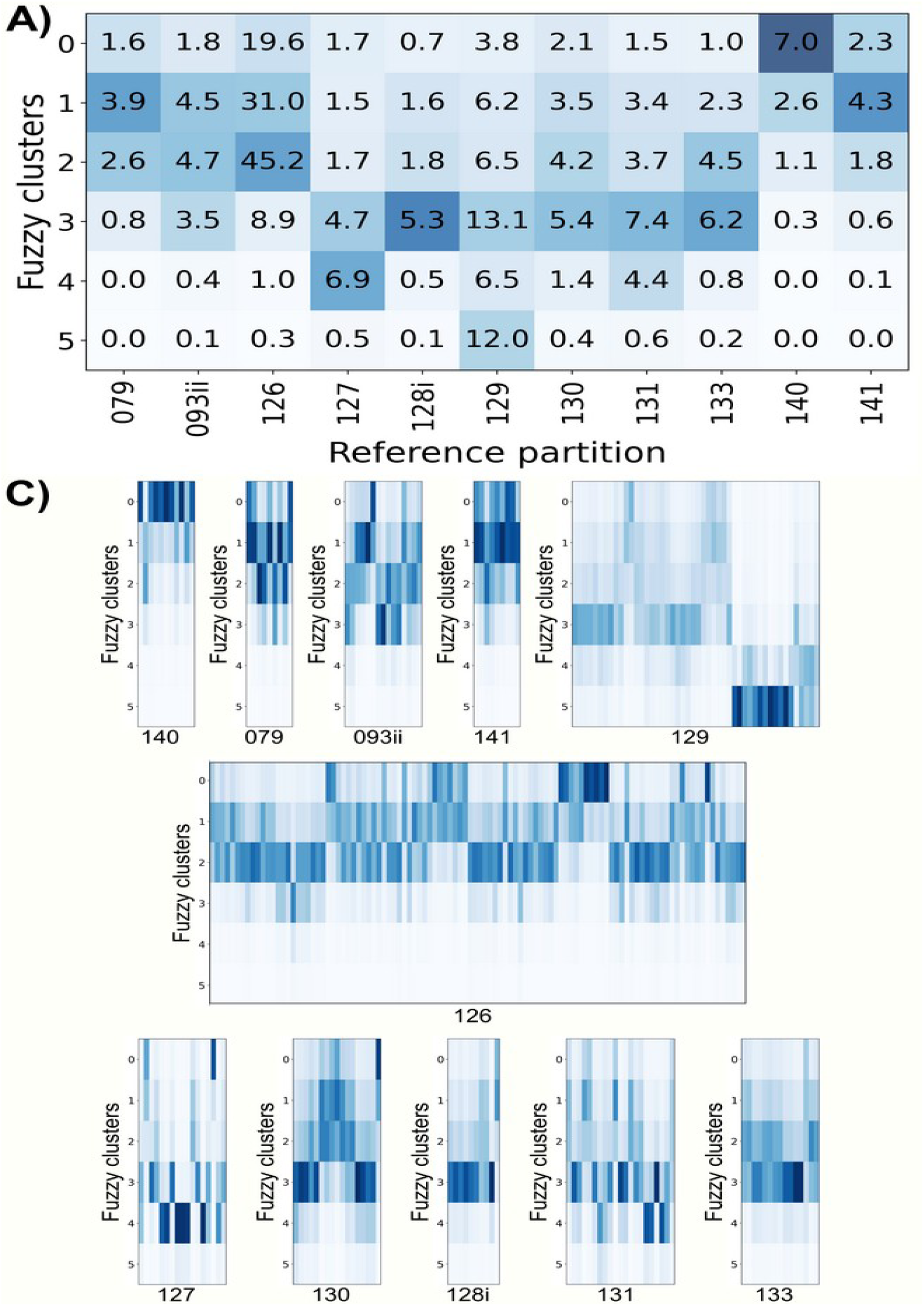
**A) Correspondence table between fuzzy clusters and catalogue call types for the six-cluster solution obtained with the “7M10S” feature set B) UMAP plot coloured by fuzzy clusters (top) or catalogue call types (bottom) C) Colour-coded membership scores of individual calls, ordered by catalogue call types.**

Some fuzzy clusters showed a direct one-to-one correspondence to a catalogue call type. Call type 140 was frequently singled out in one fuzzy cluster (Cluster 0 in Fig 9A). A subset of calls from call types 126 and 129 also showed a direct association with one or several fuzzy clusters (see Cluster 2 and 5 in Fig 9, respectively).

Many different call types showed strikingly similar patterns of membership score distribution. Most clustering solutions described the dataset as a linear continuum of calls and the distribution of call types along this continuum was always the same (Fig 9B). Call types 140 and 141 were always associated with the first fuzzy clusters and could be separated when the number of fuzzy clusters increased. Then, calls of call types 079, 093ii and 126 had similar membership scores to the next fuzzy clusters. A subset of call type 079 appeared to be strongly similar to call type 141 (Cluster 1 in Fig 9A). Calls from call types 127, 128i, 129, 130, 131, and 133 were consistently grouped in the last fuzzy clusters (Cluster 3 in Fig 9A). When the number of fuzzy clusters increased, a portion of call types 127, 129 and 131 could be separated from the rest (Cluster 4 in Fig 9A). Finally, a few samples of call type 129 were isolated from the rest of the dataset when using large feature sets (Cluster 5 in Fig 9A).

These associations of catalogue call types within fuzzy clusters reflected the structural similarity of call types. Call types 079 and 141 were both low frequency vocalisations with many harmonics and an overall level frequency, except at call onset and offset (Fig 1). The stereotyped calls reconstructed from the first fuzzy clusters, to which vocalisations of type 079 and 141 had the highest membership scores, showed a similar structure (Fig 5 and Supplementary Materials Fig S8). Call types 127, 128i, 129, 130, and 131 were often grouped together in the same fuzzy cluster. All these call types shared a similar structure: a sharp increase in frequency at call onset, followed by an asymmetrical u-shape with a rapid decrease in frequency and a slow increase until the end of the call (Fig 1). This asymmetrical u-shaped variation could be seen in some stereotyped vocalisation obtained with the finest precision reconstruction methods (“**10M7S”** and “**10M10S”** feature sets, Supplementary materials File S7).

There were also consistent differences between call types and fuzzy clusters. For instance, call type 079 was split between two fuzzy clusters with a strong contrast in membership scores between individual calls (Fig 9C). A portion of call type 129 (the rightmost vocalisations in Fig 9C) was consistently isolated from the other calls by the FC procedure. A single call from call types 127 and 128i was strongly associated with the first fuzzy cluster, while the other ones were located at the other end of the continuum (Fig 9C). Table 2 (below) presents an overview of the comparison between fuzzy clusters and catalogue call types for the different FC solutions. The absence of clear one-to-one correspondence and the description of the dataset as a continuum from call types 140 and 141 to call types 127, 131, and 129 are clearly visible in the table. For more information about the exact comparison between the FC and catalogue-based classification, you will find correspondence tables for all realisations of the FC algorithm in the Supplementary materials File S9.

**Table 2.**
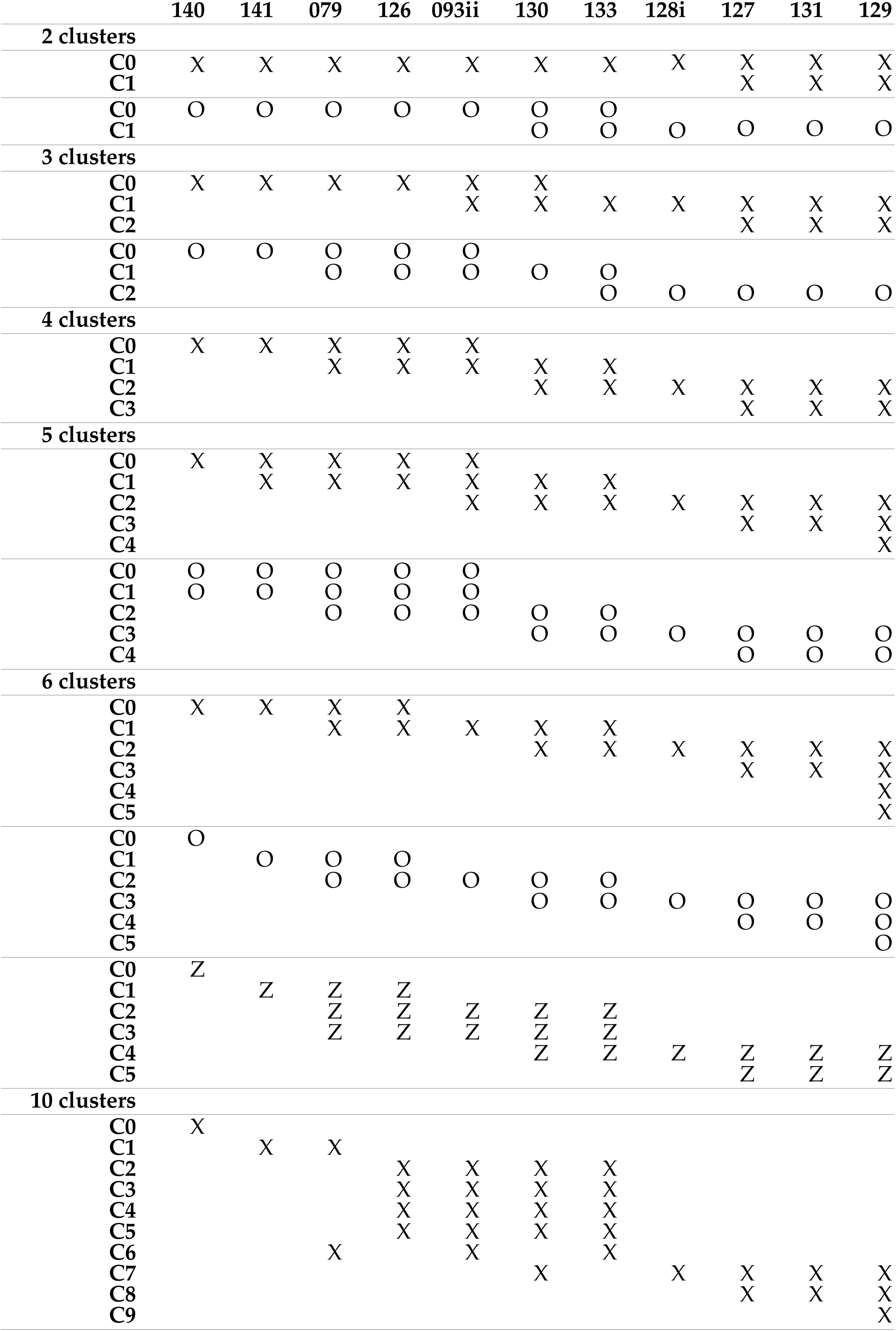
Correspondence between the fuzzy clusters and the catalogue call types for the different clustering solutions. Each column correspond to a catalogue call types, and each line to a fuzzy cluster. The symbols pictures which fuzzy clusters calls from a given call type belong to. For instance, in the two-cluster solution (top), calls of type 126 belong mostly to fuzzy cluster C0, while calls of type 127 are distributed between fuzzy clusters C0 and C1. When clustering solutions with the same number of fuzzy clusters but obtained with different feature sets differed, the alternative solutions are presented below the main one with a different symbol. 2-cluster solutions: O – **“5M5S”** solution. 3-cluster solution: O – small feature sets. 5-cluster solutions: O – small feature sets. 6-cluster solutions: X – large feature sets, O – medium-sized feature sets, Z – small feature sets. Note the overall consistency of the correspondence between fuzzy clusters and catalogue call types: call types140 and 141 always correspond to the first fuzzy clusters, and call types 127, 131, and 129 always to the last ones. Correspondence tables for all realisations of the FC algorithm can be found in the Supplementary materials File S9.

## Discussion

### Summary of our results

We implemented a novel unsupervised classification procedure, based of Mel frequency cepstral coefficients for the description of animals’ vocalisations and the fuzzy clustering algorithm for the classification of those vocalisations. We tested this classification procedure against a particularly challenging set of long-finned pilot whale calls, for which a previous operator-dependant classification was available. Our classification procedure described the dataset as an alignment of overlapping fuzzy clusters. The FC algorithm detected stable divisions of the dataset into two to ten categories, with most support for the divisions in five and six fuzzy clusters: these clustering solutions were detected using more feature sets and were stable for larger intervals of fuzziness (Fig 4 and Table 1). The different clustering solutions obtained with feature sets of different sizes and with different values of fuzziness, were highly consistent. This observation provides added weight to the detected structures in the dataset, since they appeared prominent at different spectral and temporal resolutions.

### Fuzzy clustering outcomes

Detailed analyses of the FC outcomes provided precise information about the graded nature of the vocal repertoire of long-finned pilot whales. The analysis of typicality, on the one hand, is a straightforward way to characterise individual calls as stereotyped – highly similar to the templates around which the vocal repertoire is organised – or graded – sharing similarities with several such templates. However, typicality provides no information about the location of graded calls within the vocal repertoire. Triangular gradation plots require a more thorough analysis than typicality, however they provide precise information about the position of individual calls in-between the fuzzy cluster centres. These two approaches complement each other and provide useful insight into the structure of graded vocal repertoires.

Typicality and triangular plot analyses revealed varying degrees of typicality according to fuzzy clusters. Half of the fuzzy clusters showed much overlap and were densely surrounded by calls, which characterised a very graded portion of the pilot whales’ call repertoire (Supplementary Materials Fig S6). The second half of the fuzzy clusters were farther away from each other and located is sparser regions of the dataset: they corresponded to a more stereotyped, but less frequently used portion of the vocal repertoire (Supplementary Materials Fig S6).

The FC procedure described the dataset as an alignment of fuzzy clusters. Gradation occurred almost exclusively between successive fuzzy clusters in the sequence (Figs 7-8). On the one hand, this could mean that the call repertoire of long-finned pilot whales can be interpreted as a single continuum from low frequency, slightly rising vocalisations to vocalisations dominated by a u-shaped, mid frequency element. On the other hand, this could mean that our classification procedure lacked power to describe finer structures in the vocal repertoire of long-finned pilot whales, either because our descriptive features were not precise enough or because of intrinsic characteristics of the FC algorithm. It is worth noting that the dataset we used in this study corresponded to a small portion of the vocal repertoire of the species. The alignment of fuzzy clusters was the prominent structure detected at several temporal and spectral resolutions. However, some clustering solutions deviated from this pattern and detected a subset of call type 129 as lying at the edge of the vocal repertoire. More research is needed to explore the reliability of the FC classification procedure, especially for large and graded vocal repertoires.

### Comparison to manual classification

The FC procedure did not replicate manual classification. We described between two and ten fuzzy clusters, with classifications into five and six fuzzy clusters showing the strongest stability. Therefore, the precision of the FC procedure was slightly lower than the manual classification, with 11 call types. There was a large degree of agreement between both classification procedures, with one-to-one correspondences between call types and fuzzy clusters and consistent associations of multiple call types in the same fuzzy clusters. It is worth noting that there were also discrepancies between the manual classification and the FC procedure: some call types were systematically distributed over several fuzzy clusters, with stark contrasts of membership scores between subsets of vocalisations. Similar results have been observed in previous studies (23).

Our unsupervised procedure was much faster than the audiovisual inspection of the spectrograms: the feature extraction and the classification algorithm took a few days to compute, compared to several months of analysis by human operators. Vester and colleagues ran a hard clustering algorithm on a larger set of long-finned pilot whales and could defined five broad categories (19). In a previous study, Wadewitz and colleagues used FC to classify primate vocalisations: they described five clusters of primate calls (21). Therefore, our FC procedure did not achieve lower precision with an unbalanced, highly graded set of vocalisations than could be expected from previous applications of this algorithm to bioacoustics. In addition, using soft classification did not result in fewer categories being defined than using hard classification (19).

### Dimensionality of the FC algorithm

We faced no strong issues with the high-dimensionality limitations of the FC algorithm (34,35). The FC procedure described the same structures in the dataset using feature sets of different sizes. We did not reach a clustering solution with a single fuzzy cluster using the smallest feature sets. Indeed, the smallest number of fuzzy clusters in solutions obtained with the **5M5S** feature set was five. It may be necessary to adjust the algorithm parameters, and especially **p_max_**, to the size of the feature set.

As the number of dimensions in the feature set increased, the fuzziness values in the clustering solutions of interest decreased and there were fewer realisations of the algorithms to use in further analyses. Using the largest feature set – **10M10S**, 101 features – we only detected three clustering solutions: with two, three, and six fuzzy clusters. Therefore, it appeared that the FC algorithm is safe to use with dataset of up to 100 dimensions. This prescriptive limit may change with the size of the dataset (32,33).

### Future directions for FC in bioacoustics research

Different perspectives for the development of the FC procedure would be the extension of the procedure to automated call segmentation, the improvement of the objective function, its association with playback experiments, and including an adaptation of the descriptive features to different study species.

Indeed, our procedure now includes feature extraction (taken from the “pylotwhale” python package) and classification (with the “fuzzyclustering” python package). Another time-consuming step of research in bioacoustics is call segmentation. Several automated approaches to call segmentation are available (e.g. 39,40). Including an automated call segmentation step to the FC procedure could substantially improve the time spent auditing and classifying a dataset.

Then, the current objective function of the FC procedure only relies on cluster compactness, that is how close individual samples are from the centre of the fuzzy clusters. The generalised intra-inter silhouette is an adaptation of the silhouette measure of clustering quality (41) to fuzzy clusters (42). This index takes into account both cluster compactness and cluster separation and can be adapted to both hard and soft classifications for comparison purposes. It is therefore a better measure of clustering quality than the current objective function of the FC algorithm. However, its calculation implies a large number of distance comparisons and proved to be computationally too expensive, even for our dataset of moderate size. An optimised coding of the generalised intra-inter silhouette could improve our FC procedure.

In this article, we present an unsupervised approach to describe the vocal repertoire of long-finned pilot whales using MFCC and FC. However, there is no guarantee that the fuzzy clusters we defined are relevant to the animals. Playback experiments using stereotyped and graded vocalisations in differential behavioural and social contexts are still needed to understand how long-finned pilot whales use their rich and flexible vocal repertoires. Associated with the FC approach to account for call variability, they may shed a new light on the communicative functions of long-finned pilot whale vocalisations.

The Mel scale we used in this article is well-suited for the study of Mammals, such as long-finned pilot whales, as it mimics the logarithmic scale of pitch perception by mammalian ears. We could adapt the FC procedure to different study species by taking into account available information about the hearing of these species. This adaptation could take the form of alternative frequency scales for non-mammalian species, or the adaptation of the maximum levels and boundaries of the Mel filter to the hearing threshold of the study species at different frequencies (see 17 for an example with elephant hearing).

### Concluding remarks

In conclusion, we developed an unsupervised and fast classifier which takes into account and quantifies the graded nature of animal vocalisations. Testing against a set of long-finned pilot whale calls, our procedure achieved a precision similar to hard clustering approaches and slightly lower than manual classification, with the added-value gradation characterisation. Our procedure can readily be adapted to other study species and may provide a more nuanced description or vocal repertoires beyond the graded-stereotyped dichotomy. The graded nature of vocalisations can have a biological significance for animals (e.g. graded variation with context of emission 10,or graded encoding of aggressive intent 43).We strongly encourage future studies to adopt soft classification procedure such as ours in order to take a quantitative approach to the study of gradation in the vocal repertoires or animals.

## Supporting information captions

**File S1. ZIP archive containing spectrograms and Mel spectra of individuals long-finned pilot whale calls in the dataset.** Spectrograms were computed using 1024 points segments, a Hanning window, and 50% overlap. Mel spectra were computed using 40 Mel filters and summarised over ten equal time slices. The file name indicates the catalogue call type the vocalisation was assigned to.

**File S2. ZIP archive containing PCA plots for all realisations of the FC algorithm selected for further analyses.** The file names give the feature set used (in the form “XMYS” for **X** MFCC and **Y** time slices), the number of fuzzy clusters (in the form “Zc” for **Z** clusters) and the fuzziness value (“p224**”** means **p** = 2,24).

**File S3. ZIP archive containing UMAP plots for all realisations of the FC algorithm selected for further analyses.** The file names give the feature set used (in the form “XMYS” for **X** MFCC and **Y** time slices), the number of fuzzy clusters (in the form “Zc” for **Z** clusters) and the fuzziness value (“p224**”** means **p** = 2,24).

**File S4. ZIP archive containing all triangular gradation plots.** Individual triangular plots showing gradation between a single pair of fuzzy clusters and compound figures showing all pairs of fuzzy clusters for a given FC realisation. The file names give the feature set used (in the form “XMYS” for **X** MFCC and **Y** time slices), the number of fuzzy clusters (in the form “Zc” for **Z** clusters), the fuzziness value (“p224**”** means **p** = 2,24), and identify which pair of clusters is shown for individual plots (“cx-cy”, with **x** and **y** the number of the clusters).

**Fig S5. Reconstruction of the centre of the fuzzy clusters for the six-cluster solution obtained using the “5M7S” feature set.** Each line corresponds to one FC realisation. The realisations are ordered from top to bottom by increasing fuzziness. The order of the clusters, from left to right, is the same throughout the manuscript. Note the low spectral resolution of stereotypes reconstructed from **“5M”** feature sets: two bands are visible, centred on the 11^th^-12^th^ and the 25^th^ Mel filters (0.8-1.3 kHz and 5-6 kHz) and spanning several Mel filters.

**Fig S6. Reconstruction of the centre of the fuzzy clusters for the six-cluster solution obtained using the “10M10S” feature set.** Each line corresponds to a realisation of the FC algorithm. The realisations are ordered from top to bottom by increasing value of fuzziness. The spectral resolution of the reconstructed calls was higher when using ten MFCC than when using five (Fig S5). Here, we reconstructed more distinct frequency bands, and they spanned a smaller number of Mel filters. Note the overall similarity of the reconstructed vocalisations with Fig S5.

**File S7. ZIP archive containing all stereotyped calls reconstructed from the coordinates of the fuzzy cluster centres.** Individual reconstructed stereotypes or compound figures showing all reconstructed stereotypes for a given clustering solution. The file names give the feature set used (in the form “XMYS” for **X** MFCC and **Y** time slices) and the fuzziness (“p224**”** means **p** = 2,24). For compound plots, the file names also give the number of fuzzy clusters (in the form “Zclusters” for **Z** clusters). For individual reconstructions, they identify the reconstructed cluster centre (as “clusteraonb”, **a** being the number of the cluster, and **b** the total number of fuzzy clusters).

**Fig S8. Reconstructed cluster centres for the different clustering solutions obtained with the “10M5S” feature set A) Two-cluster solution B) Three-cluster solution C) Four-cluster solution D) Five-cluster solution E) Eight-cluster solution.** A single example is shown for each clustering solution, drawn approximately from the centre of the fuzziness interval. Note the similarity of the cluster centres across clustering solutions. Note also the similar continuum in panels A-D. Panel E shows a clustering solution that derived slowly from the general pattern: the dataset is described here as a linear continuum between Cluster 0 and Cluster 6. Cluster 7 is located outside of this alignment of fuzzy clusters.

**File S9. ZIP archive containing correspondence matrices between fuzzy clusters and catalogue call types for all realisations of the FC algorithm.** The table represent either the correspondence for a single realisation of the FC algorithm or the correspondence for a clustering solutions using average membership scores (identified by “global” in the file name). The file names also give the feature set used (in the form “XMYS” for **X** MFCC and **Y** time slices), the number of clusters (“zc” for **z** clusters), and the fuzziness (“p224**”** means **p** = 2,24).

**File S10. ZIP archive containing typicality histograms, coloured by fuzzy clusters, for all realisations of the FC algorithm.** Typicality histograms are presented alone or associated with a PCA or UMAP plot for the visualisations of the fuzzy clusters. The file names give the feature set used (in the form “XMYS” for **X** MFCC and **Y** time slices), the number of clusters (“zc” for **z** clusters), and the fuzziness (“p224**”** means **p** = 2,24).

